# Evolution of behavioral flexibility and the forming and breaking of habits

**DOI:** 10.1101/2025.08.29.673035

**Authors:** Olof Leimar, Sasha R. X. Dall, Peter Hammerstein, Alasdair I. Houston, Bram Kuijper, John M. McNamara

## Abstract

The formation of habits, whereby learnt actions come to be performed automatically with repetition and practice, is a well-established focus for studies in psychology. This contrasts with evolutionarily motivated studies of learning, which typically view behavior as either learnt or fixed, to elucidate the ecological conditions where each predominates. Here, we envisage habit formation functioning to free up limited mental resources (e.g., attention), potentially improving an individual’s ability to multitask. As an ecologically relevant case, we investigate exploration-exploitation in foraging under predation risk. In our model, a forager does not know the quality of feeding options, but can learn from the rewards they give. When the environment occasionally changes, individuals can attend to exploration of feeding options in the new conditions. The options can then become habitually exploited, freeing attention for antipredatory vigilance. Via evolutionary simulations, we show that evolutionarily stable forming and breaking of foraging habits can substantially reduce mortality from predation, without drastically reducing foraging success, when environmental conditions remain stable enough between changes. We identify factors promoting habit formation, such as the repetition of actions that yield predictable rewards, and also discuss the role of habits more generally, including their relation to ideas about bounded rationality in theories of decision making. In conclusion, we argue that exploration-exploitation strategies involving the forming and breaking of habits are a type of behavioral flexibility that is likely to be selected for under a range of ecological conditions.

**Significance statement:** The forming and breaking of habits has long been a focus of study in psychology, as well as appearing in many discussions about human behavior. Surprisingly, the topic has been virtually absent from evolutionary studies of learning in non-human animals, despite featuring prominently in Darwin’s discussion of the evolution of species-specific behaviors. Our aim is to change this, by theoretically investigating a possible evolutionary explanation for habits, namely that habits enhance an individual’s ability to multitask. We model the combination of foraging and antipredator vigilance, which is important for many animal species in their natural environments. We conclude that the forming and breaking of habits can promote behavioral flexibility, helping animals to achieve important tasks in different environments.

---

Habits, as studied in experimental psychology, are modes of behavior where individuals come to perform actions as more-or-less automatic responses, triggered by stimuli or circumstances they repeatedly encounter. The formation of habits is typically guided by goal-directed learning processes [1, 2, 3, 4, 5], making it a common feature of human task learning. Indeed, habit learning mechanisms seem to be conserved across the mammalian species that have been subject to laboratory reinforcement learning research [6] and are suggested to be shared by all vertebrates [7, 8]. The continuum between learnt, habitual and fixed behavior suggests a more nuanced view of learnt behavior than is typical for evolutionarily motivated studies of learning. Despite early ideas about the inheritance of habits for the evolution of species-specific behaviors [9], behavioral ecologists typically consider the evolutionary stability of learning strategies compared to innate, fixed behavioral responses [10, 11]. Here we suggest that the forming and breaking of habits represent important types of behavioral flexibility and deserve to be taken into account in evolutionary investigations of animal behavior.

An important property of habits is that, in terms of mental resources, they are efficient ways of performing tasks, by letting actions be guided reflexively by stimuli and contexts [5]. In contrast to non-habitual behavior, which involves continuous learning and updating, habitual task performance enables individuals to free cognitive resources like attention to devote to other concerns [3, 5]. It is generally recognized that limited attention can constrain multitasking [12, 13]. Here we use evolutionary simulations to show that habit-enhanced multitasking can be selectively favored, by letting animals efficiently combine foraging and vigilance against predators. Our work is a further development of the study of foraging-vigilance trade-offs, which has long been a central topic in behavioral ecology [14, 15, 16].

To analyze the forming and breaking of habits, we use the idea of a trade-off between exploration and exploitation. This trade-off figures prominently in the fields of psychology, behavioral ecology, and machine learning [17, 18, 19, 20, 21, 22], but there is no consensus about the cognitive mechanisms that control such behavior. Our approach is to assume that transitions between exploration and exploitation correspond to the forming of new habits and the breaking of old habits [1, 2, 5, 6, 23].

We examine situations where the reward distributions of different feeding options are unknown but remain constant over a period of time, after which they can change. In nature, a change could for instance be due to a new season or moving to a new location, and there could be several such environmental changes over an individual’s lifetime. If each successive environment is constant for long enough, it would be advantageous to first learn about the feeding options and then to become habitual in foraging, thus balancing the attentional demands of foraging and vigilance. Such situations could favor the evolution of psychological mechanisms for the forming and breaking of habits. It is our aim to establish the conditions under which these mechanisms are advantageous, by examining cases where exploration and learning influence predation risk, as well as cases with little or no influence. We also investigate factors influencing the speed of habit formation and illustrate situations where habits should not form.

In addition to changes in rewards, individuals can perceive other cues indicating new environments. These cues could be signs of seasonal change, or for an individual to have arrived to a new location. There are psychological studies investigating the effects of cues of changes of context [3, 4, 5, 24, 25]. In our modelling, we assume that such additional cues of change are available.

The elements of our models include psychological mechanisms of exploration and learning, as well as rules for the forming and breaking of habits. For exploration, we use traditional action-value learning, but modified such that the initial estimated values of options can be high [20]. We refer to the approach as optimistic exploration (OE). This approach is well established in theoretical work on reinforcement learning [20, 26, 27, 28, 29, 30, 31] and has the advantage of a close correspondence to traditional assumptions about learning in experimental psychology. For comparison, in the supplements we also show results with so-called Thompson sampling (TS) [32, 33] as the exploration strategy. TS is a Bayesian statistical approach that is argued to show good performance in some situations [33], but it is not known if it is used by real animals. Both OE and TS originated in the study of multi-armed bandits [20, 33].

To implement habit formation, we use the well-established idea in cognitive modelling [2, 34] that habits are characterized by consistency of choice, i.e. the tendency to choose a given action in a given situation. In our simulations, we express the consistency of choice as a running estimate of the probability to repeat the most recent action. We assume that as an individual’s estimate of its consistency of choice passes a threshold, the individual switches to habitual foraging, at the same time increasing its vigilance against predators. For the reverse transition from habitual foraging to exploration, we assume that individuals can evolve to respond to a cue of a new environment, by resetting the attention to learning about foraging options, including restarting optimistic exploration.

In the following, we illustrate the kinds of environmental volatility we investigate and show results from individual-based evolutionary simulations of the performance of exploration-exploitation strategies. We find that the forming and breaking of foraging habits has the potential to significantly reduce mortality from predation, without drastically reducing foraging success. We therefore argue that there is likely to be substantial natural selection for this form of behavioral flexibility.

## Results

Figure 1 shows a situation that could promote the forming and breaking of habits. A forager is faced with a number of feeding options (we use 8 feeding options). Environments are randomly generated, in the sense that the mean reward for each option is drawn from a log-normal distribution (Fig. 1A). In each of a number of foraging trials, there is a choice of feeding option to use in that trial. Each feeding option delivers positive, log-normally distributed rewards, with a certain mean and (log-scale) random variation around this mean (Fig. 1B). In our analyses we assess performance over two or more successive environments. Ideally, in such situations a forager should explore to find the highest yielding option in an environment. The intuition behind optimistic exploration (OE) is sketched in Fig. 1C. By starting with high initial estimates of reward rates, which then are reduced through the sampling of options, a forager is incentivized to sample all options and, eventually, to focus on the highest yielding one. If a sufficiently thorough exploration can be achieved early in the period of an environment, and foraging then becomes habitual, the cost of increased predation risk from reduced vigilance during exploration and learning can be fairly small (Fig. 1D).

**Figure 1:**
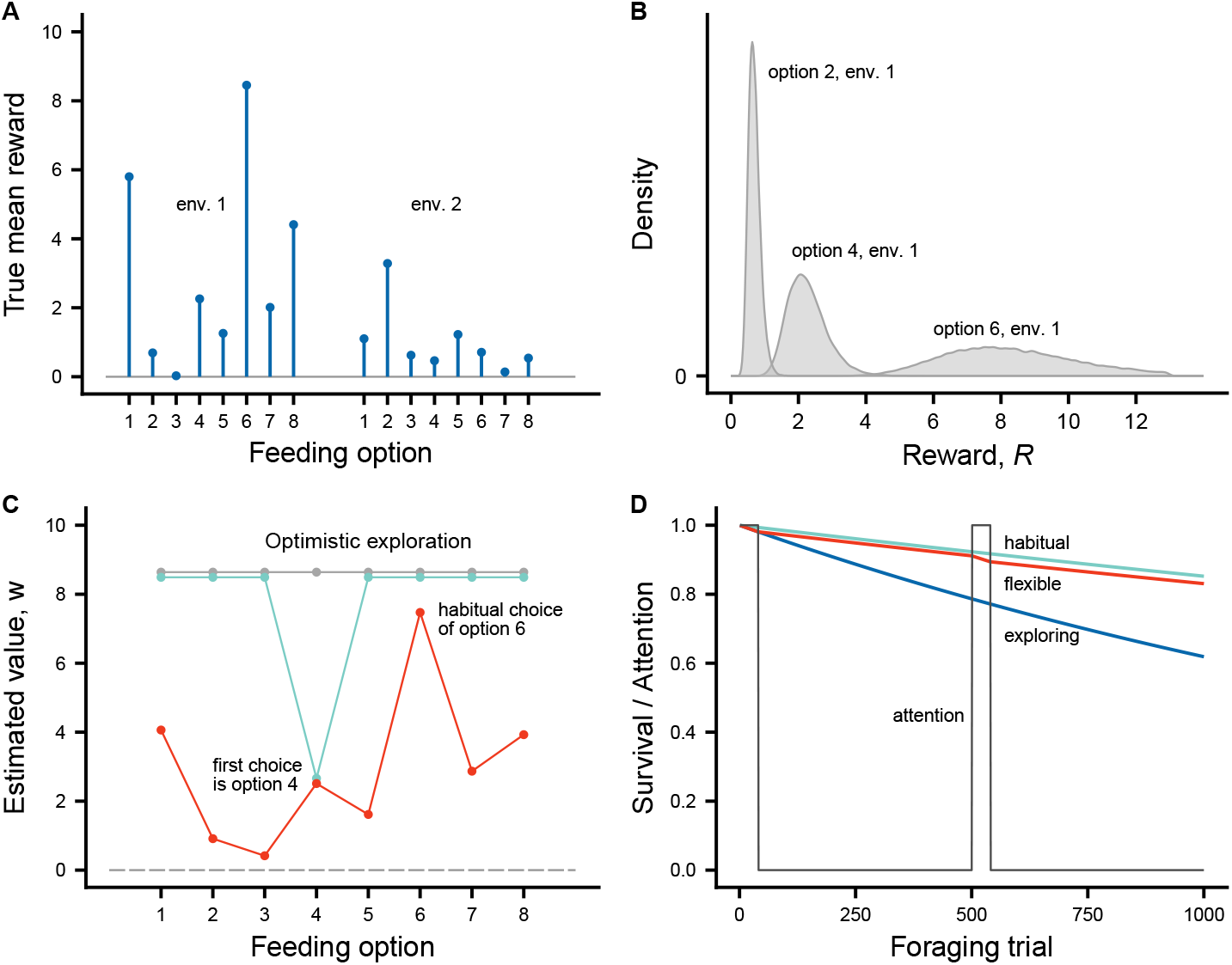
Illustration of model elements. Individuals make and break foraging habits, thereby trading off vigilance and exploration of food sources. As an example, there are two successive environments, each with 500 foraging trials, resulting in a maximal foraging lifetime of 1000 trials. In each environment, there are eight feeding options, or stimulus types. The options vary in reward distributions between environments. By devoting attention to the exploration of feeding options individuals can increase their feeding rates, at a cost of decreased vigilance. **(A)** Mean reward values for the feeding options in two successive environments. Each mean value is independently drawn from a log-normal distribution with a log-scale mean of *ρ*_0_ = 0 and log-scale SD of *σ*_0_ = 1. **(B)** Examples of the distribution of rewards for three different options, taken from the options available in environment 1. The reward from a given option is log-normally distributed with a given mean and a log-scale SD of *σ*_*R*_ = 0.25. **(C)** Illustration of a learning strategy of optimistic exploration. The curves show an individual’s estimate of the values of options (used for soft-max choice) in environment 1 at different points in time. Estimates start high (gray curve). The first choice is option 4, after which estimates are given by the greenish curve. The estimated values after all options are explored and the individual’s behavior is habitual are given by the red curve (curves are shifted slightly to avoid overlap). **(D)** Survival for full habitual foraging (full vigilance; greenish curve; 0.85 at trial 1000), full attention to exploration (blue curve; 0.62 at trial 1000), and an intermediate case of initially exploring each environment and then becoming habitual (red curve). The gray curve shows a corresponding switching on and off of attention for a flexible strategy.

Our approach is to run individual-based simulations in which foragers have evolving traits, including an initial value estimate (*w*_0_), a value learning rate (*α*_*w*_), a choice consistency update rate (*α*_*h*_), a habit-formation consistency threshold (*ĥ*), and a softmax choice inverse temperature parameter (*β*), as detailed in methods. Figure 2 shows the evolutionary equilibrium outcome for a situation as in Fig. 1. The regret values in Fig. 2A,C are theoretical quantities (deviations of expected reward of actual choice from the expected reward of an optimal choice), and are not observed by foraging individuals but are instead often used to evaluate the performance of exploration-exploitation strategies [33, 35]. The red curve in Fig. 2A shows that the regret is very close to zero during trials with habitual foraging, while the survival stays close to a maximal full-vigilance value. This illustrates the potential efficiency of the forming and breaking of habits. The example of a particular individual (Fig. 2C,D) illustrates exploration in an environment where one option (number 3) is considerably more rewarding than the others. After 12 trials of exploration, the individual switches to habitual foraging.

**Figure 2:**
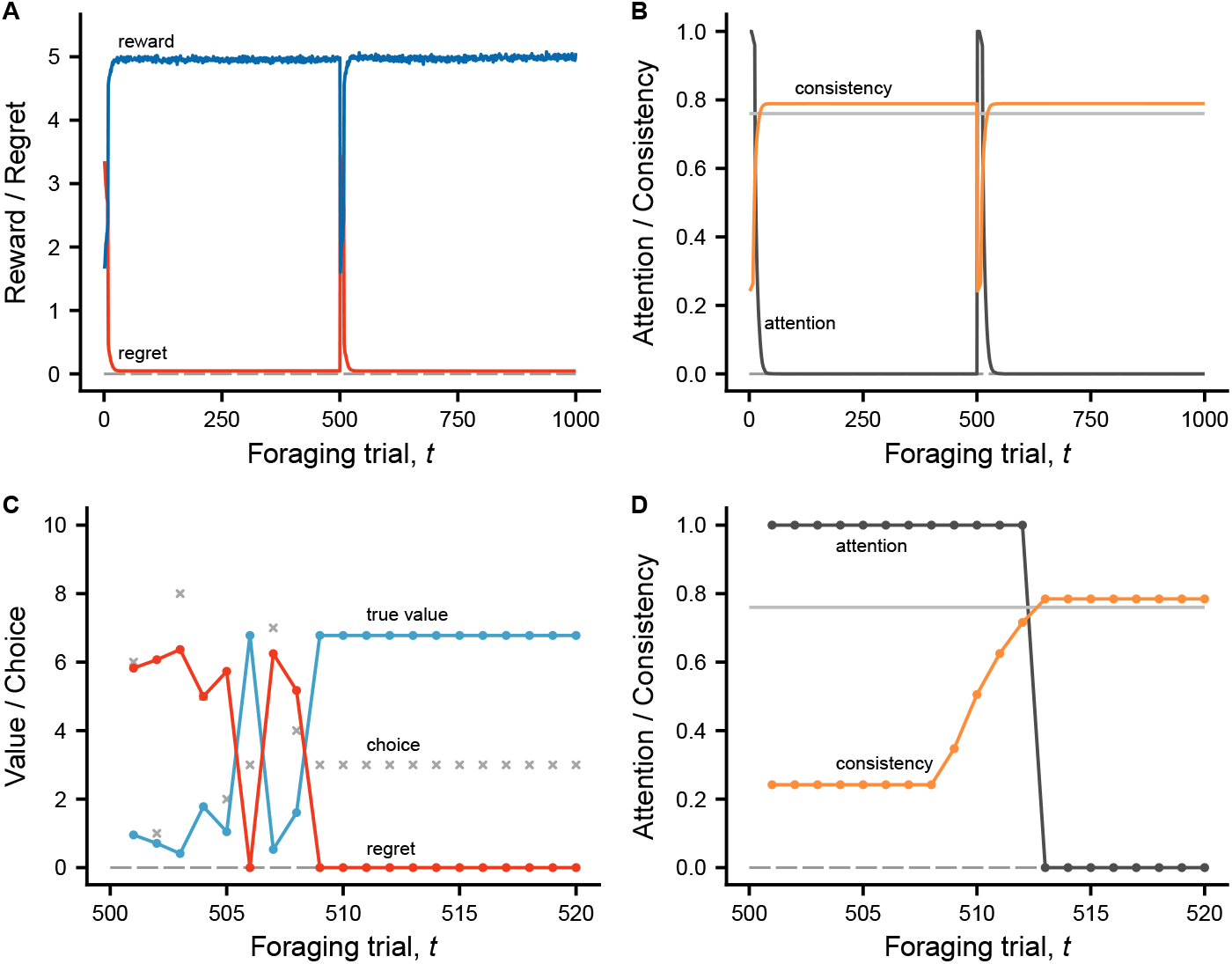
The forming and breaking of habits at an evolutionary equilibrium, for a situation as illustrated in Fig. 1. In each of two successive environments, there are eight feeding options, with expected reward values drawn from a log-normal distribution (log-scale mean *ρ*_0_ = 0 and SD *σ*_0_ = 1) and trial-to-trial random reward variation as in Fig. 1B (*σ*_*R*_ = 0.25). At the start of each environment, individuals observe a reliable cue, indicating that the environment has changed. There are a total of 4000 simulated individuals, with traits for optimistic exploration and habit formation at evolutionary equilibrium. Panel **(A)** shows the mean over all individuals of the reward in each trial (blue curve), together with the mean regret (red curve). An individual’s regret in a trial measures how much the individual lost by not choosing the option with the maximum expected reward (zero regret indicates an optimal choice). The mean attention to foraging (dark gray) and the mean consistency of choice (orange) are in panel **(B)**. Individuals turn off attention to foraging and become habitual when consistency goes above an evolved threshold *ĥ*, indicated by a gray horizontal line, after which the update of consistency stops. The consistency measures the degree to which the current choice is the same as the most recent choice. Panels **(C)** and **(D)** show an example of one individual, for the first 20 trials of the second environment. The successive choices are indicated by gray crosses in **(C)**, together with the corresponding expected (true) values *w* (blue-green) and regret values (red). The average survival over 1000 trials for the habit forming individuals is around 0.84, close to the maximal value for full vigilance, and better than the expected survival of 0.62 for full attention to exploration. Trait values: *w*_0_ = 9.34, *α*_*w*_ = 0.90, *α*_*h*_ = 0.24, *ĥ* = 0.76, *β* = 15.5.

By modifying the amount of randomness in rewards, we can test the observation from psychology that habits are formed more readily when rewards are more predictable [2, 5]. Our evolutionary model provides support for this idea. As illustrated in Fig. 3, the time until habit formation in a new environment is about three times longer when rewards are relatively noisy, compared to a case of little random reward variation around the mean value of options.

**Figure 3:**
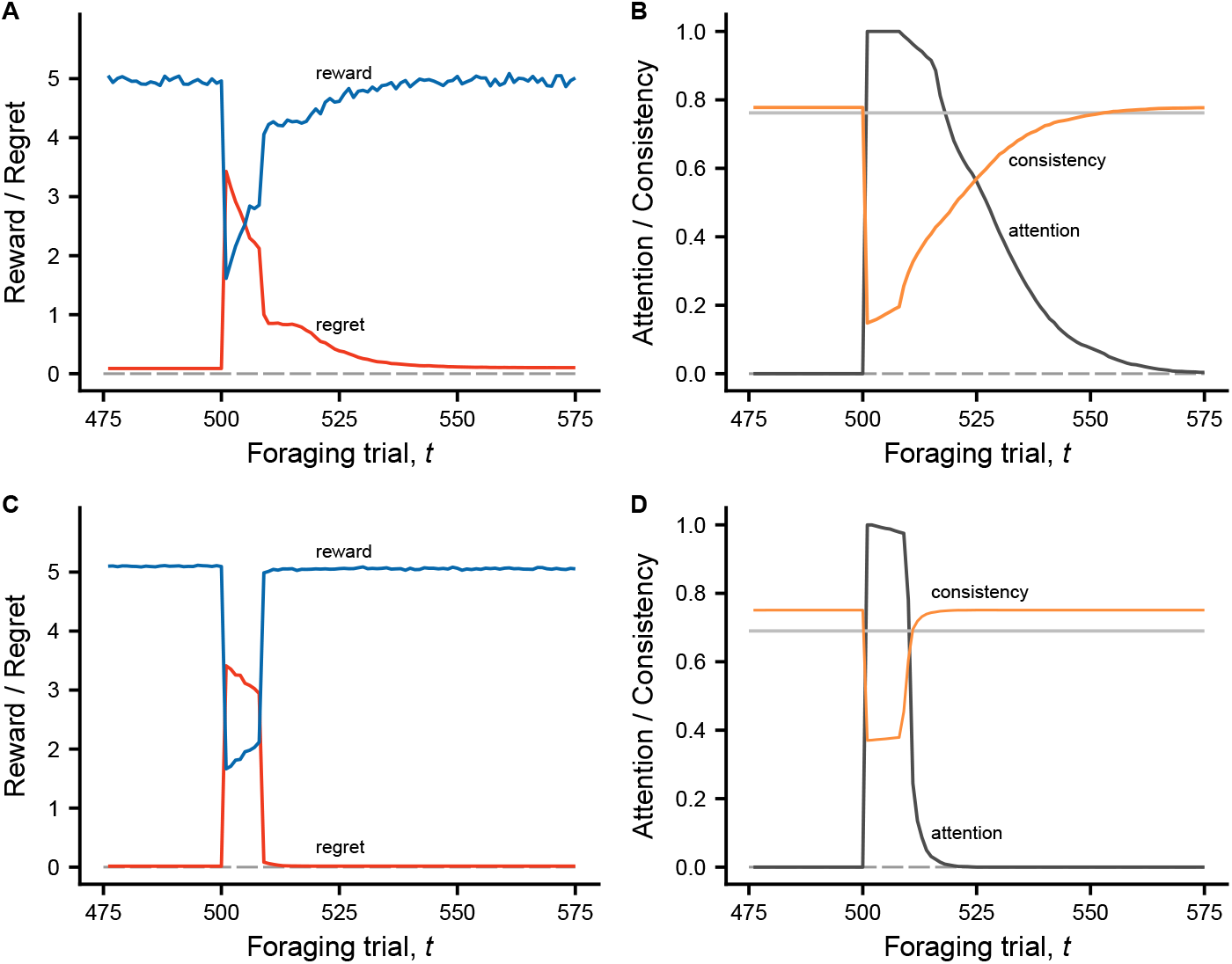
The influence of the predictability of rewards on the speed of habit formation. Illustration of means of reward, regret, attention, and choice consistency for evolutionary equilibria of traits of optimistic exploration and habit formation. The situations are similar to Fig. 2, except that the random variation of rewards for a given option is high (*σ*_*R*_ = 0.50) in panels **(A)** and **(B)**, and low (*σ*_*R*_ = 0.10) in panels **(C)** and **(D)**. The panels show trials around the time of environmental change at *t* = 500 trials. With less predictable rewards **(A, B)** habitual foraging takes longer to develop compared to the case with more predictable rewards **(C, D)**. A factor behind the difference is that the consistency update rate *α*_*h*_ is lower in **(A, B)** than in **(C, D)**. At evolutionary equilibrium, the trait values for lower, higher predictability are *w*_0_ = 5.76, 13.62, *α*_*w*_ = 0.64, 0.97, *α*_*h*_ = 0.15, 0.37, *ĥ* = 0.76, 0.69, *β* = 12.6, 17.0.

The evolutionary simulations for Fig. 3 illustrate a notable property of the exploration strategy we use here (OE). When observations are noisy, in the sense that random variation (*σ*_*R*_) is comparable to (or even larger than) variation among mean rewards of feeding options, the initial estimate of reward (*w*_0_) evolves to a smaller value; *w*_0_ *≈* 6 in Fig. 3A,B and *w*_0_ *≈* 14 in Fig. 3C,D. This means that the term ‘optimistic exploration’ is appropriate mainly for observations with relatively little noise.

Intuition might suggest that if exploration and learning do not influence predation risk, habit formation should not evolve, but that is not what our analyses show. As an illustration, the situation in Fig. 4 is the same as in Fig. 3, except that attention to foraging does not influence predation risk. The outcome is that the forming and breaking of habits still evolve, although the time until habit formation in a new environment is twice as long or more in Fig. 4 compared to Fig. 3. Another difference is that the fitness benefits of habit formation are substantially greater when there are effects of predation risk than without such effects (see supplements on fitness effects of habit formation). The explanation for habit formation in Fig. 4 might be that, after a certain amount of sampling of options, and given the finite time horizon, if one is doing well in terms of current rewards it does not pay to continue sampling.

**Figure 4:**
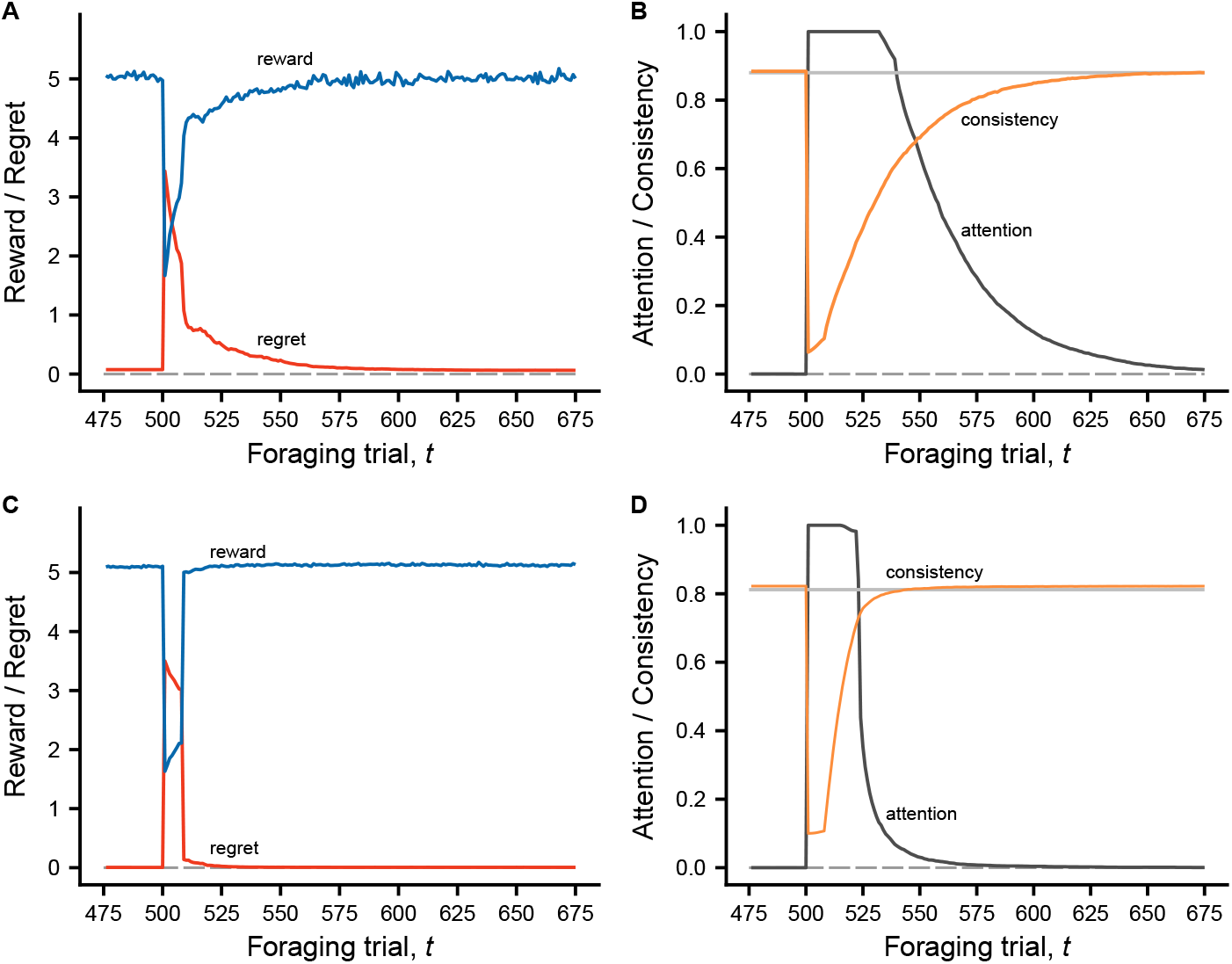
The forming and breaking of habits can also be selectively favored when exploration and learning do not influence predation risk. The reward distributions of options are the same as in Fig. 3, but the mortality schedules differ. Here, the attention to exploration and learning does not influence predation risk. The panels show means of reward, regret, attention, and choice consistency for evolutionary equilibria of traits of optimistic exploration and habit formation, for two cases differing in the predictability of rewards. The random variation of rewards for a given option in high (*σ*_*R*_ = 0.50) in panels **(A)** and **(B)**, and low (*σ*_*R*_ = 0.10) in panels **(C)** and **(D)**. The panels show trials around the time of environmental change at *t* = 500 trials. With less predictable rewards **(A, B)** habitual foraging takes longer to develop compared to the case with more predictable rewards **(C, D)**, and each of these times is longer than is the case in Fig. 3 (note that the scales on the x-axes differ between Figs. 3 and 4). At evolutionary equilibrium, the trait values for lower, higher predictability are *w*_0_ = 4.62, 13.63, *α*_*w*_ = 0.62, 0.96,, *α*_*h*_ = 0.06, 0.10, *ĥ* = 0.88, 0.81, *β* = 15.8, 16.2.

This raises the question of when habit formation should be selectively disadvantageous. Among the various possibilities, our analyses identify high environmental volatility, in the sense of more rapidly changing environments, in combination with little of no influence of exploration and learning on predation risk, as disfavoring the formation of habits (see Fig. S1 in supplements, which also illustrates the effects of less reliable cues of environmental change).

In the supplements, we also show results corresponding to Figs. 2, 3, 4, but using Thompson sampling instead of optimistic exploration (Figs. S2, S3, S4). Overall, our results support a general conclusion about selection for the forming and breaking of habits: a clear separation among the reward distributions of available feeding options (i.e., predictable rewards), together with readily available habit-breaking indicators of new environments, and with environments staying relatively constant for an appreciable period, act to promote the evolution of habit-mediated exploration-exploitation strategies and promote rapid habit formation in new environments.

## Discussion

Based on our evolutionary analyses, we propose that the forming and breaking of habits is an important type of behavioral flexibility. The flexibility is advantageous over a range of situations, including when there are wholesale environmental changes. Our evolutionary perspective corroborates current views on habits in psychology [3, 4, 5]. An example of a point of agreement is that habit formation is known to be promoted by periods of environmental stability, and also by the repetition of actions that yield predictable rewards [2, 5], which is in accordance with our results in Fig. 3. We also found that habits form more quickly when the benefits of multitasking are greater (comparing Figs. 3 and 4), which is intuitive but not previously studied.

In psychological studies of learning, changing environments are often discussed in terms of volatility of rewards [36], with reversal of rewards perhaps being the most studied [2, 37, 38], together with the case of continual shifting of the reward values or the probabilities of reward [17, 39]. These forms of volatility correspond to aspects of environments encountered in nature, and could matter for habitual responses to specific stimuli or situations. In our work here, the main focus is on the forming and breaking of habits as a consequence of wholesale environmental changes.

Different meanings have been given to the term behavioral flexibility [40, 41, 42], with different experimental measures or tests of such flexibility. Reversal learning is perhaps the most commonly used test, and it is in this general sense we are suggesting that the forming and breaking of habits is a form of behavioral flexibility. The specific habit-forming and -breaking mechanisms assumed in our model are likely to be simplifications of real mechanisms. Thus, habits may form gradually [2, 5] rather than through a threshold mechanism. In our model, the breaking of habits depends on specific cues, in accordance with work on the effects of context changes [3, 4, 5]. An alternative possibility, inspired by work in neuroscience on detecting and reacting to unexpected changes in the world [18, 43, 44], would be that a lowering of the rate of rewards below a threshold causes a transition from habitual foraging to exploration.

Habit formation in our model is a process of first exploring options and then switching to habitual foraging. This is qualitatively similar to a strategy of so-called ‘satisficing’, as developed in the study of bounded rationality [45, 46, 47, 48]. In theories of economic decision making, a satisficing strategy searches through alternatives until a threshold for acceptability is met, at which time searching stops. In making this comparison, one should note that evolutionary equilibria in our model represent optimal collection of rewards under the constraints of predation risk and a limited time period, where optimality is over the space of strategies given by our trait space [49]. This differs from the current usage of the term satisficing in theories of decision making: optimization under constraints is regarded as distinct from bounded rationality [50, 51]. It then seems that our analysis of the forming and breaking of habits is closely related to but distinct from ideas of bounded rationality in economics. In any case, habit formation is likely to play an important role in human decision making.

There is a wide field of investigation into mechanisms of learning. The switching on and off of learning in our model can be seen as a version of the much-studied idea that learning rates should respond to environmental characteristics, for instance with higher learning rates in more volatile environments [52, 53, 54, 55]. Further, the concept of attention has long been used to develop ideas about learning [56, 57, 58, 59, 60, 61]. Our model assumption of a simple switching on and off of attention to exploration of feeding options should be seen as a first-step simplification of possibly complex dynamics, involving gradation and shifts between different reward-predicting stimulus dimensions. Even so, there is an empirically well-supported tradition in psychology and neuroscience of so-called dual-process models, with essentially discrete modes of behavior [1, 2, 5, 62, 63, 64]. The exploratory and habitual behaviors in our model are meant to represent such modes.

In addition to these models, there could to be other ways of being efficient in combining foraging and vigilance. One example is a model where individuals can commit to foraging over a period of several trials, while at the same time being sensitive to indirect, predator-related cues that can interrupt the foraging [65]. Individuals can then forage while also maintaining a certain vigilance, by being sensitive to cues, such as rustling sounds, indicating the presence of a predator. This need not be related to habit formation, but it is also possible that habitual foraging allows an individual to direct more attention to a range of cues that can give information about predation risk. In general, exploration, foraging, and vigilance are tasks with several demands on mental capacities. For instance, exploration in nature could involve paying attention to and remembering locations and surroundings of different feeding options, in this way showing complexities beyond responding to a multi-armed bandit.

The essential role played by habitual foraging in our model is to enhance multitasking, through improved vigilance with only minor changes in foraging success. The combination of foraging and vigilance is sometimes studied in psychology [65, 66] and, as mentioned, has long been of interest in behavioral ecology [14, 15]. This interest includes the role of attention in detecting predators [12, 13, 67] and the importance of vigilance for survival [68]. There are experimental approaches to measuring variation in vigilance, for instance comparing the time until detection of a (model) predator between different kinds of aggressive behavior in prey animals [69], or a similar comparison for fish prey surrounded by either familiar or unfamiliar conspecifics [70]. Such approaches, involving measuring the reaction time to the presentation of a model predator, could serve to diagnose whether or not a foraging habit has been formed. This would then be an addition to current experimental methods of diagnosing habitual behavior in psychological research [3, 4, 5].

A characteristic property of habits is of course the tendency to use a particular action in a given situation. This could be used to diagnose and thus establish a time scale for the shift from exploration to exploitation of a foraging individual. For habit formation to be advantageous in nature, this time scale should be shorter than the time scale of significant environmental change. In addition to seasonal changes and new locations, one needs to also take into account other changes, such as those deriving from foraging itself. The latter will be particularly relevant for group-living animals with frequency-dependence in the rewards of foraging.

In addition to foraging, there are other ecologically significant activities that sometimes need to be combined with antipredatory vigilance, for instance social behavior, like aggression, caring for offspring, and attracting mates. The formation of habits for these activities might then be advantageous. Further, if developmental plasticity from early learning leads to variation in consistent behavioral traits such as boldness and aggression [71], habit formation might contribute to animal personality variation. A general conclusion is then that further study of habits should be worthwhile for evolutionary and behavioral ecology, including through modelling, field observation, and ecologically-motivated experiments.

## Methods

Here we give an overview of the models we analyze, including the type of environments we study and the strategies individuals use in these environments. The strategies have parameters, and some of these parameters are assumed to be genetically determined traits. The evolution of these traits is studied in individual-based simulations, where the reproductive success is taken to be proportional to the total amount of reward an individual accumulates over its lifetime, up to death by predation or, otherwise, over the total number of trials assumed for the model. See Tables S1, S2 for equilibrium trait values and Table S3 for notation.

### Distribution of rewards from feeding options

The mean values 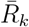 of options vary between environments (see for instance Fig. 1A). Each mean value 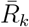 is independently drawn from a log-normal distribution (ensuring positive values) with a log-scale mean *μ*_0_ and log-scale SD *σ*_0_. In the examples in the main text *μ*_0_ = 0 and *σ*_0_ = 1. As is illustrated in Fig. 1B, there is also random variation in the rewards from a particular feeding option. We assume that this variation is log-normally distributed. Thus,

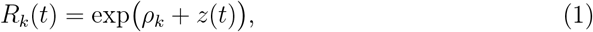

where the *z*(*t*) are independent and normally distributed with mean zero and standard deviation *σ*_*R*_ and with *ρ*_*k*_ such that 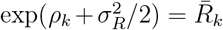, where 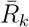 is the mean reward from the option. This gives 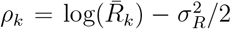. As illustrated in Fig. 1B, we use *σ*_*R*_ = 0.25 in many of our examples. We sometimes use the notation *R* to mean *R*_*k*_(*t*) for the chosen option *k* in trial *t*.

### Predation risk and attention dynamics

In principle, individuals could pay attention to and learn about certain stimulus dimensions or features while ignoring others, but as a simplification we consider all learning about feeding options as regulated in an on/off fashion. Attention is indicated by a 0/1 variable *a*, with *a*(*t*) = 1 in trial *t* if the individual is exploring and learning in that trial (using either of the two approaches described below). The rate of predation in trial *t*, as a function of attention, is then

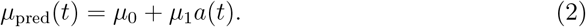

For our analyses we have used *μ*_0_ = 0.00016 and *μ*_1_ = 0.00032 (except for cases of no influence of attention on predation risk, where we used *μ*_1_ = 0.0). Over a total of 1000 trials, this gives a survival of 0.62 with full attention to exploration and 0.85 for pure habitual foraging. This means that habit formation allows a forager to decrease predation mortality by at most around 60% of its value for full attention to exploration.

For deciding when to turn off attention and become habitual, we assume that while learning an individual keeps track of its consistency of choice *h*_*k*_ for each feeding option, as follows:

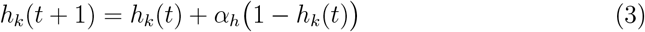

for the chosen option *k* in trial *t* and

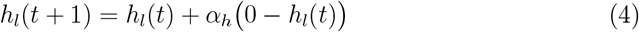

for each non-chosen option *l* ≠ *k*. The *h*_*k*_ can be thought of as measures of ‘habit strength’, satisfying 0 ≤ *h*_*k*_ *<* 1. We sometimes write consistency of choice to mean the consistency *h*_*k*_ of the chosen option on a given trial. This form of consistency of choice has been used previously [2] and has also been referred to as a ‘choice kernel’ [34]. In our model, an individual turns off attention to exploration in trial *t* if

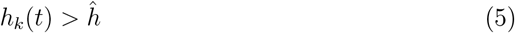

for the chosen option *k*, in which case *k* becomes the habitual choice. In order to allow evolution towards no habit format, we allow values *ĥ >* 1.

### Cues of environmental change

We assume that, on the first trial of a potentially new environment, an individual observes an environmental cue *z*_e_, indicating the probability that the environment has changed. For the results in the main text, we use ‘reliable cues’, i.e. cues that indicate a probability close to one. We assume that *z*_e_ is normally distributed with mean *μ*_e_ and standard deviation *σ*_e_, and a relation between the cue value and the probability of environmental change as in logistic regression, such that

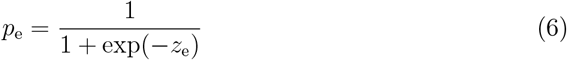

is the probability of a change. Individuals use a threshold *ê* to determine if they should restart exploration, either OE or TS, and break any current habit, resetting if

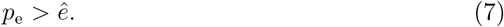

The parameters *ĥ* and *ê*, influencing the forming and breaking of habits, are implemented as genetically determined traits in our evolutionary simulations.

### Optimistic exploration (OE)

We use action value learning [20] for the exploration part of the strategy, with the added element that the starting estimated values of the feeding options can be relatively high (‘optimistic initial values’; [20]). This is essentially a version of classical Rescorla-Wagner learning [72], adapted to operant conditioning.

So if *w*_*k*_(*t*) is an individual’s estimate of the reward value of choosing option *k* in trial *t*, the starting value is assumed to be *w*_*k*_(0) = *w*_0_. Ideally, for optimistic exploration, *w*_0_ should be higher than any of the actual mean reward values of options in the current environment (cf. Fig. 1C). If option *k* is chosen in trial *t*, the individual updates its estimate to *w*_*k*_(*t* + 1), where

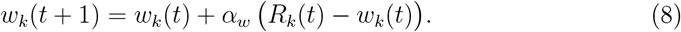

Here *α*_*w*_ is the learning rate, for simplicity assumed to be constant in time and the same for all options, and *R*_*k*_(*t*) is the reward perceived by the individual from interacting with a stimulus of type *k* at time *t*. The quantity

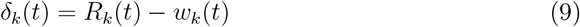

is referred to as the prediction error and is the difference between the reward *R*_*k*_(*t*) currently experienced by the individual and its prior estimate *w*_*k*_(*t*) of the reward. The change in the estimated feature value *w*_*k*_ in equation (8) is thus the learning rate *α*_*w*_ times the prediction error, and tends to move the estimate towards a ‘true value’.

### Soft-max choice

In each trial *t*, there is a choice between *n*_s_ = 8 feeding options, or stimulus types. An individual chooses option *k* with probability

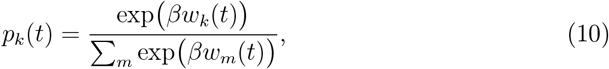

where *β* is referred to as an ‘inverse temperature’ parameter. This procedure is typically referred to as a soft-max rule, going from estimated values to a probability of choice, and it is commonly used in reinforcement learning models [20].

### Genetically determined traits

For evolutionary analysis, we assume that certain of the learning parameters are evolving traits. For optimistic exploration, the initial value *w*_0_, the learning rate *α*_*w*_, and the inverse temperature parameter *β* are such traits. In addition to these, there are also traits *α*_*h*_, *ĥ*, and *ê* for the forming and breaking of habits, as described above.

### Thompson sampling (TS)

Thompson sampling [32] is a Bayesian statistical approach with near optimal or optimal exploration-exploitation performance in certain situations [33], but it is not known if it is approximated by cognitive mechanisms in real animals. We use this form of exploration for some of the results in the Supplementary Information, to give a comparison to the OE approach in the main text.

As mentioned, we assume that the different feeding options have log-normal reward distributions. So if option *k, k* = 1, …, *K*, is chosen in trial *t*, we assume that the reward *R*_*k*_(*t*) is drawn from a log-normal distribution with parameters *ρ*_*k*_ and *σ*_*R*_, i.e., *y*_*k*_(*t*) = log(*R*_*k*_(*t*)) is normal with mean *ρ*_*k*_ and standard deviation *σ*_*R*_. For convenience we sometimes use the precision 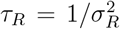 as parameter instead of *σ*_*R*_. In order to have analytic expressions for the posterior distributions, in some of our examples we further assume as prior that the parameters *ρ*_*k*_ of the options are (independently) drawn from a normal distribution with parameters *ρ*_0_ and 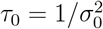. Having observed *y*_*ki*_, *i* = 1, …, *n*_*k*_ or, as a sufficient summary of observations, 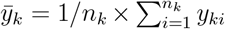, the posterior distribution for *ρ*_*k*_,

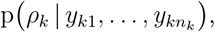

is then normal with mean

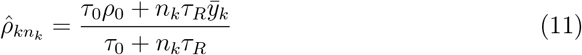

and precision

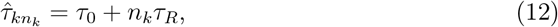

with 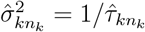 . This posterior is then used for the sampling of parameters.

### The sampling and action choice procedure

The Thompson sampling procedure is, for each *k*, to ‘draw’ parameters *ŷ*_*k*_ from the normal distributions with parameters from equations (11, 12), and then to choose the option with the largest posterior mean reward, i.e., with the largest value of

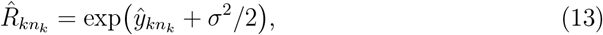

which is the same as choosing the option with the largest value of 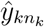 .

### Habitual foraging

For both OE and TS foragers, we assume that habitual foraging means that the last choice before the transition is repeated. The consistency of choice *h*_*k*_(*t*) stays constant during habitual foraging, and the attention variable *a*(*t*) in equation (2) is equal to zero, leading to increased vigilance.

### Order of events in a trial

In a trial, the following occurs: (*i*) check for environmental cue indicating habit breaking and resetting of exploration; (*ii*) risk of predation, following equation (2); (*iii*) choice of feeding option; (*iv*) reward; (*v*) if attention to exploration is on, update consistency of choice; (*vi*) for optimistic exploration with attention on, perform learning updates; (*vii*) if attention to exploration is on, check if it should be turned off.

### Evolutionary simulations

A simulated population has non-overlapping generations. For computational efficiency (multi-threading) in evolutionary simulations, the population was split into groups of size 10, with 400 groups, making up a total of 4000 individuals. Individuals are sexually reproducing, with one diploid locus for each trait. For simplicity, we assume that individuals are hermaphrodites, mating is random across the entire population, and the different loci are unlinked. Also for simplicity, we assume that reproductive success accumulates proportional to collected rewards over all trials (so that an individual dying from predation still has reproductive success from its rewards collected until death; other assumptions about the life history, such as discrete reproductive seasons, are of course possible).

Evolutionary equilibrium trait values were obtained by first running simulations over several 100,000 generations, recording the mean trait values every 1000 generations, until there were no systematic changes. Equilibria were then determined by running simulations for another 100,000 generations, computing mean trait values from values recorded every 1000 generations. The data for the figures were then obtained by simulating a single generation with these mean trait values, randomly generating environments for each of 4000 individuals, in order to achieve maximum sampling of environmental variation.

Simulations were implemented as C++ programs, with separate programs for OE and TS. The genetically determined traits are *w*_0_, *α*_*w*_, *α*_*h*_, *ĥ, β, ê* for OE, and *α*_*h*_, *ĥ, ê* for TS. We implemented mutation as potentially occurring for each allele of a newly formed individual, with probability 0.002 and normally distributed mutational increments. The standard deviation of the increments was adjusted to the range of variation of a given trait.

## Code availability

C++ source code for the individual-based simulations is available at GitHub, together with instructions for compilation on a Linux operating system (https://github.com/oleimar/habits2)

## Acknowledgements

We thank Redouan Bshary, Peter Dayan, and Alex Thornton for helpful comments on a previous version of the manuscript. S.R.X.D. was supported by a Royal Society Leverhulme Trust Senior Research Fellowship,

## Author contributions

O.L., S.R.X.D., A.I.H., and J.M.M. designed research; O.L. performed research; O.L. wrote the original draft; O.L., S.R.X.D., P.H., A.I.H., B.K., and J.M.M. discussed results and contributed to the editing of the final manuscript.

## Supplementary information

### Illustration of effects of higher environmental volatility

In order to identify situations where habit formation does not evolve, here we examine cases where there can be many environmental changes during an individual’s lifetime (up to 50 environments), with each environment potentially having a short duration of only 20 trials. If each such environment is drastically different, this might be an unrealistically high volatility, so we also examine cases where successive environments are fairly similar, in the sense of small values of the parameter *σ*_0_. We combine a range of cases with different values of *σ*_0_, with *ρ*_0_ adjusted to keep the overall mean rewards constant, with cases of different influence of attention to foraging on predation risk, having either *μ*_1_ = 0.00032 or *μ*_1_ = 0.0 in equation (2).

Our model assumes that individuals can observe a cue that can indicate a change of environment, or context. In our examples in the main text, these cues have been highly accurate in predicting change, but it is also of interest to investigate the effect of cues of lower accuracy. For high volatility, it could be realistic that cues indicating change are of intermediate reliability. We implement this by having a quantitative cue and letting the cue strength indicate the probability that the environment has changed, as in equation (6) in the main text. If this probability is small and the different environments are relatively similar, it might be best for habitual foragers to ignore the cue, thus avoiding predation costs of resetting attention to exploration.

These possible effects, together with the influence of exploration and learning on predation risk (parameter *μ*_1_), and the magnitude of change between environments (parameter *σ*_0_), are illustrated in Fig. S1. A main result is that habit formation is not selected for if there is a combination of high environmental volatility and no influence of exploration and learning on predation risk.

**Figure S1:**
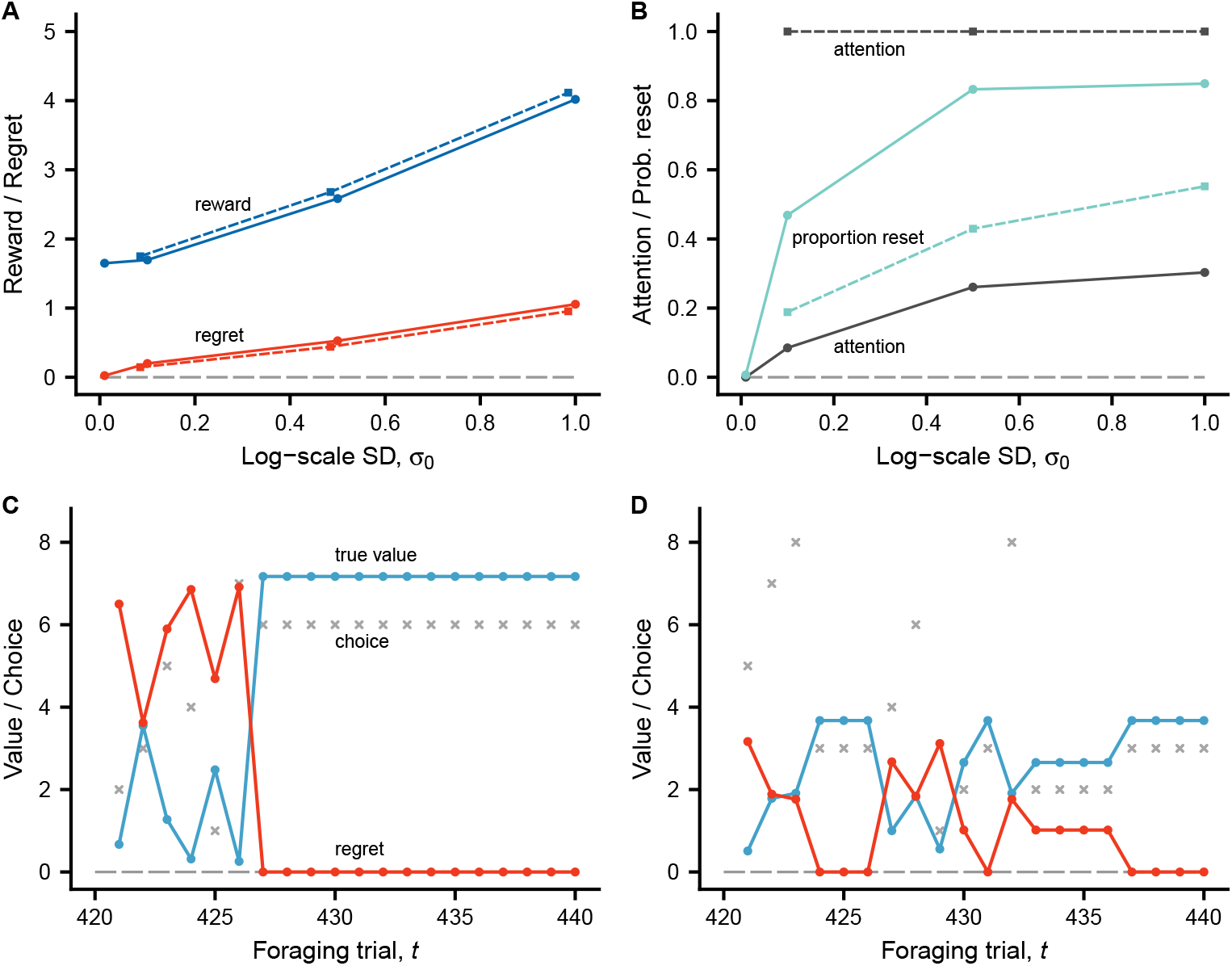
Illustration of the overall means over all individuals and trials (where the individual is alive) of reward, regret, attention, and the probability that an individual resets exploration in response to a cue, for seven cases with different parameter values *σ*_0_ and *μ*_1_. The values of *σ*_0_ are on the x-axes of panels **(A)** and **(B)**, and solid/round vs. dashed/square symbols indicate *μ*_1_ = 0.00032 vs. *μ*_1_ = 0.0. Note in particular that for *μ*_1_ = 0.0 in panel **(B)**, the evolutionary equilibrium is full attention to exploration and learning, and thus no habit formation, whereas there is forming and breaking of habits for *μ*_1_ = 0.00032. Panel **(C)** illustrates a simulated individual from the *σ*_0_ = 1, *μ*_1_ = 0.00032 case initiating habitual foraging during a brief span of a new environment lasting 20 trials. Panel **(D)** illustrates a simulated individual from the *σ*_0_ = 1, *μ*_1_ = 0.0 case correctly avoiding to respond to a cue of change.

### Comparing optimistic exploration with Thompson sampling

Figures S2, S3, and S4 show results that correspond to Figs. 2, 3, and 4 in the main text, with OE replaced by TS as the exploration strategy. Comparing Figs. 2 and 3 with S2 and S3, one sees that in these situations there is qualitative similarity between OE and TS in how habits are formed, the main difference being that the period of exploration before habit formation is longer for TS than for OE. A likely explanation for this is that OE is initially faster than TS in picking more rewarding options, whereas in the long run TS is more thorough in its exploration.

There is a qualitative difference when comparing Figs. 4 and S4, where exploration does not influence predation risk. For these environments, OE leads to the evolution of habit formation, whereas habit formation is selected against with TS. In terms of rewards collected there is, however, not a big difference, with the overall mean regrets being small and of similar magnitudes OE and TS.

With less variation in expected rewards between options and more stochastic variation in rewards, there is again qualitative similarity between OE and TS when there is no predation risk, in that there is selection for habit formation both for OE and TS, as is illustrated in Fig. S5.

### Fitness effects of habit formation

In our model, an individual’s lifetime reproductive success (LRS) is proportional to the lifetime sum of rewards obtained by the individual, up until the individual dies from predation or until the end of the maximum life span. Comparing the mean of this measure, for the examples in Figs. 3 and 4 in the main text, with simulations where individuals are constrained not to form habits (not shown in figures), we find that there is reduction of LRS from preventing habit formation of around 14% in Fig. 3, but only around 1% in Fig. 4. This illustrates that the fitness benefits of habit formation can be substantially greater when there are effects of attention to exploration on predation risk, compared to when there there are no such effects.

This stronger selection for habit formation if habitual foraging reduces predation risk is further illustrated in Fig. S6. Starting evolutionary simulations from a point of no habit formation, with predation risk there is fairly rapid evolution of habit formation (Fig. S6C,D). Also, as illustrated by the variation in the population average thresholds *ĥ* (Fig. S6A,C), with at least some habit formation, stabilizing selection towards an evolutionary equilibrium of *ĥ* is comparatively weaker.

**Figure S2:**
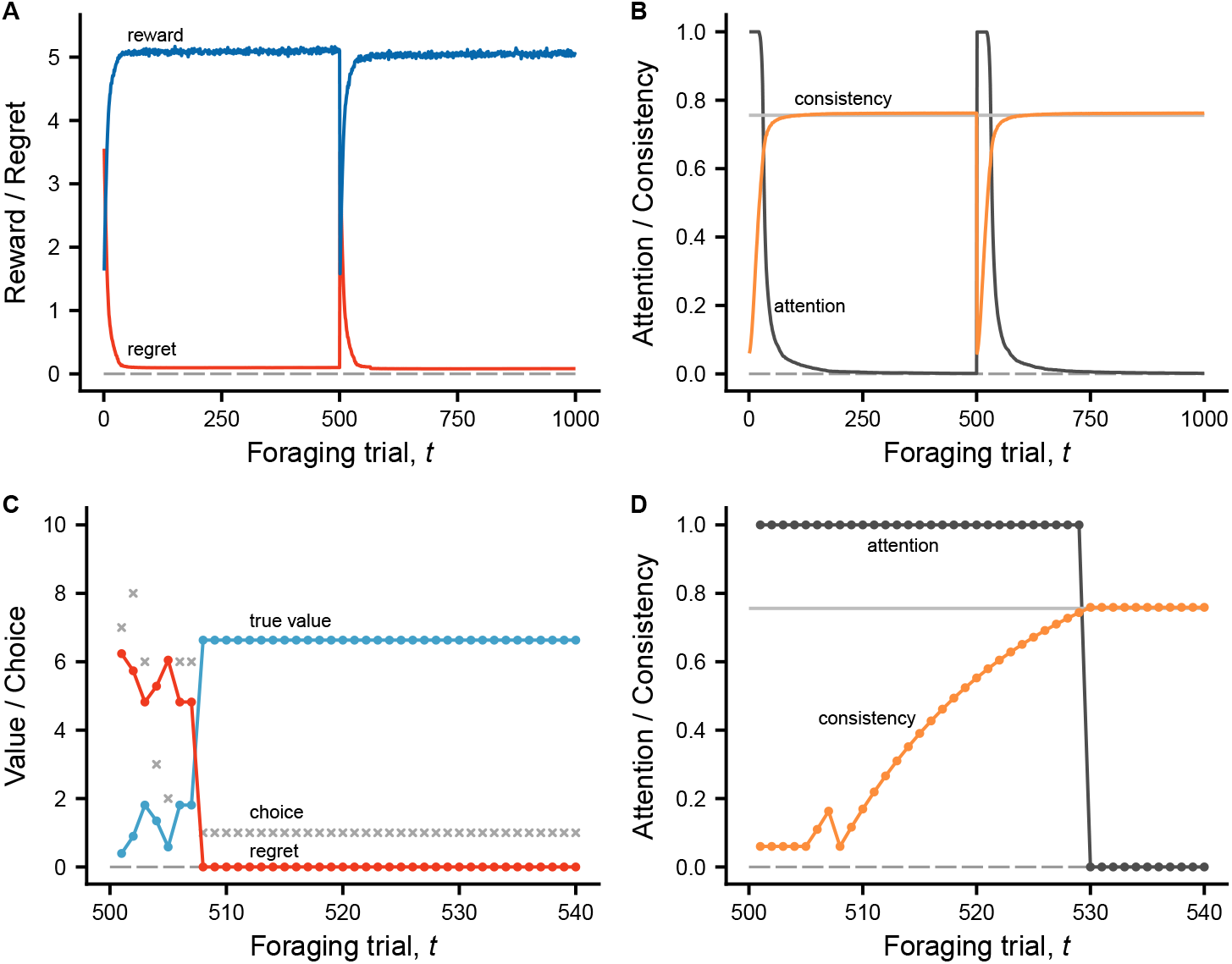
The forming and breaking of habits at an evolutionary equilibrium, corresponding to Fig. 2, but with TS instead of OE. In each of two successive environments, there are eight feeding options, with expected reward values drawn from a log-normal distribution (log-scale mean *ρ*_0_ = 0 and SD *σ*_0_ = 1) and trial-to-trial random reward variation as in Fig. 1B (*σ*_*R*_ = 0.25). At the start of each environment, individuals observe a cue, indicating that the environment has changed. There are a total of 4000 simulated individuals, with traits for habit formation at evolutionary equilibrium. Panel **(A)** shows the mean over all individuals of the reward in each trial (blue curve), together with the mean regret (red curve). The mean attention to foraging (dark gray) and the mean consistency of choice (orange) are in panel **(B)**. Individuals turn off attention to foraging and become habitual when consistency goes above an evolved threshold *ĥ*, indicated by a gray horizontal line, after which the update of consistency stops. The consistency measures the degree to which the current choice is the same as the most recent choice. Panels **(C)** and **(D)** show an example of one individual, for the first 40 trials of the second environment. The successive choices are indicated by gray crosses in **(C)**, together with the corresponding expected (true) values *w* (blue-green) and regret values (red). The average survival over 1000 trials for the habit forming individuals is around 0.83, slightly less than in Fig. 2, but better than the expected survival of 0.62 for full attention to exploration. Trait values: *α*_*h*_ = 0.06, *ĥ* = 0.76.

**Figure S3:**
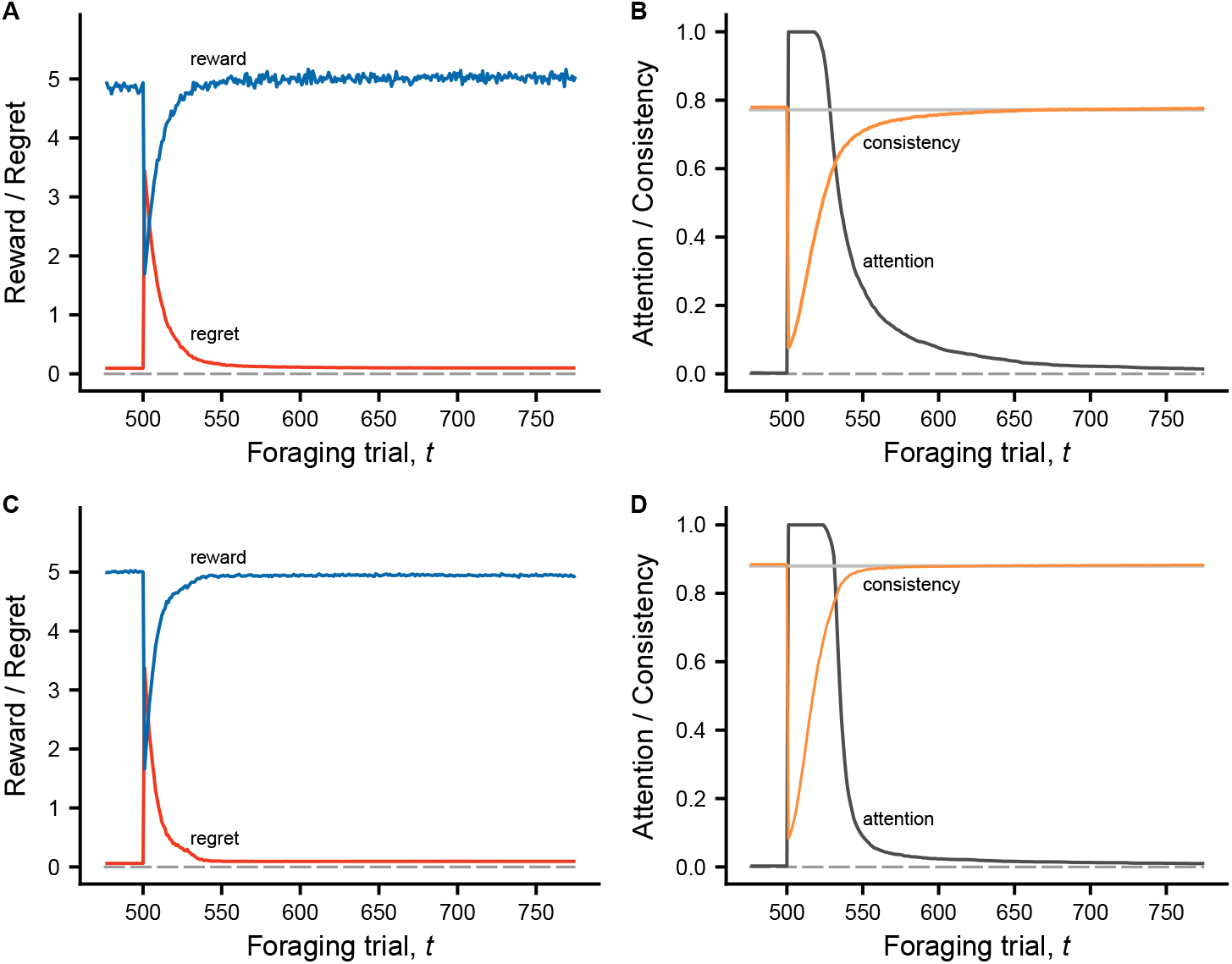
The influence of the predictability of rewards on the speed of habit formation, corresponding to Fig. 3, but with TS instead of OE. Illustration of means of reward, regret, attention, and choice consistency for evolutionary equilibria of traits of habit formation. The situations are similar to Fig. S2, except that the random variation of rewards for a given option is high (*σ*_*R*_ = 0.50) in panels **(A)** and **(B)**, and low (*σ*_*R*_ = 0.10) in panels **(C)** and **(D)**. The panels show trials around the time of environmental change at *t* = 500 trials. With less predictable rewards **(A, B)** habitual foraging takes longer to develop compared to the case with more predictable rewards **(C, D)**. A factor behind the difference appears to be that TS entails less repetition of choice in **(A, B)** than in **(C, D)**, resulting in a slower buildup of consistency of choice *h*_*k*_(*t*). At evolutionary equilibrium, the trait values for lower, higher predictability are *α*_*h*_ = 0.08, 0.08, *ĥ* = 0.77, 0.88.

**Figure S4:**
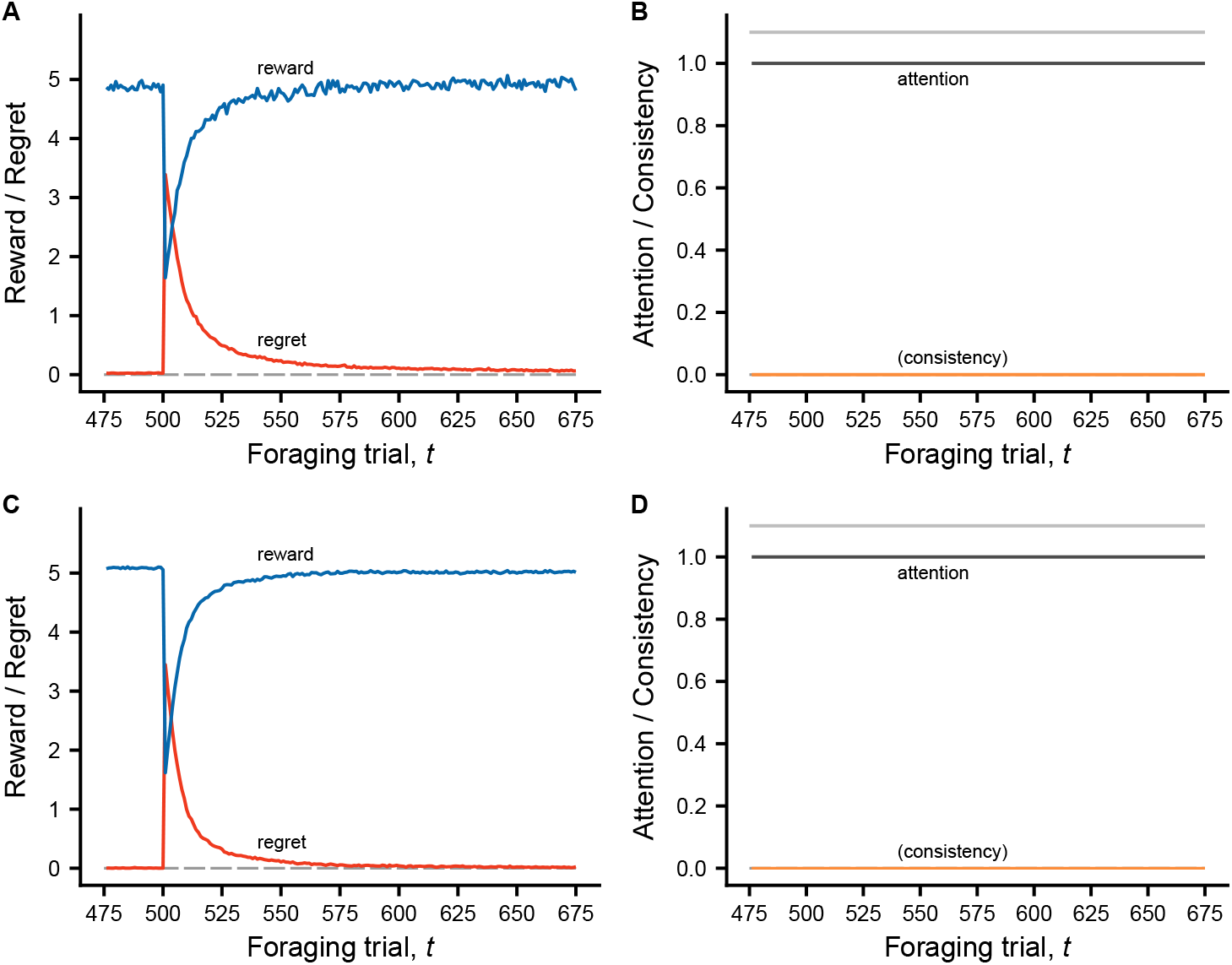
For TS and with environmental volatility/stochasticity as here, the forming and breaking of habits is selected against when exploration and learning do not influence predation risk. For our model, habit formation can be prevented if the habit-formation consistency threshold *ĥ* evolves to a value greater than one, which happens in these cases. The reward distributions of options are the same as in Fig. S3, but here the attention to exploration and learning does not influence predation risk. The panels show means of reward, regret, and attention for TS. Because the trait *ĥ* evolved to *ĥ >* 1 in both these cases, choice consistency is irrelevant for habit formation and attention. The random variation of rewards for a given option is high (*σ*_*R*_ = 0.50) in panels **(A)** and **(B)**, and low (*σ* _*R*_ = 0.10) in panels **(C)** and **(D)**. The panels show trials around the time of environmental change at *t* = 500 trials.

**Figure S5:**
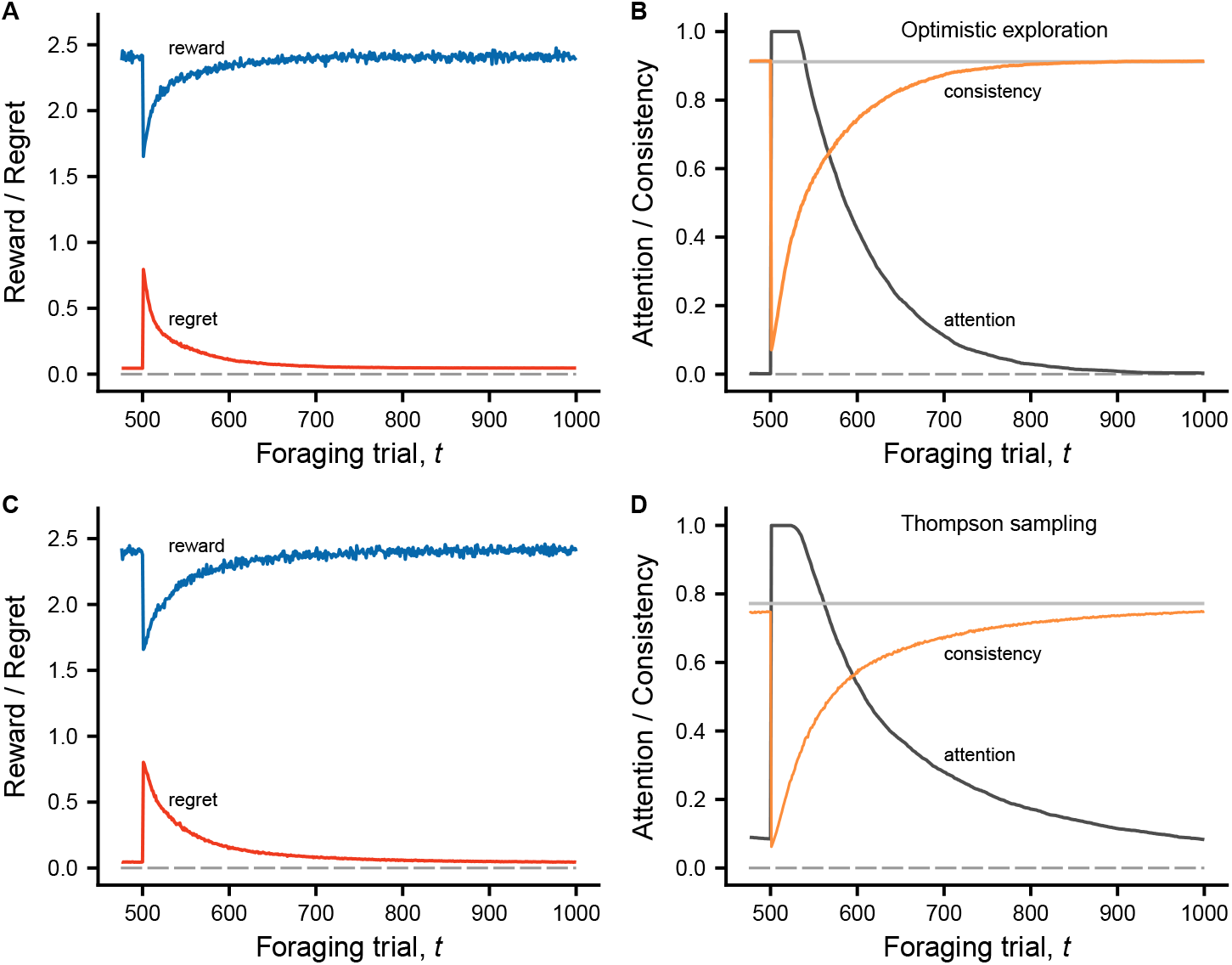
Comparison of exploration and habit formation between OE, in **(A)** and **(B)**, and TS, in **(C)** and **(D)**, for a case with fairly little variation between options in expected rewards, but considerable reward stochasticity, resulting in low predictability. The parameters of the reward distributions are *ρ*_0_ = 0.455, *σ*_0_ = 0.30, and *σ*_*R*_ = 0.5. There are two successive environments, each of 500 trials, without any mortality (*μ*_0_ = *μ*_1_ = 0), so there is no predation advantage of habitual foraging. Panels **(A, C)** show mean reward and regret, and **(B, D)** show attention to exploration and consistency of choice, for 4000 simulated individuals, over the trials in the second environment. Times until habit formation are longer than for the examples in the main text and in Figs. S2, S3, S4. For TS the time until habit formation is very variable among individuals, with a fraction of the individuals continuing to explore over the entire duration of the environment, as seen in **(D)**. The trait values are at an evolutionary equilibrium, with *w*_0_ = 1.93, *α*_*w*_ = 0.20, *α*_*h*_ = 0.07, *ĥ* = 0.91, *β* = 12.4 for OE and *α*_*h*_ = 0.06, *ĥ* = 0.77 for TS.

**Figure S6:**
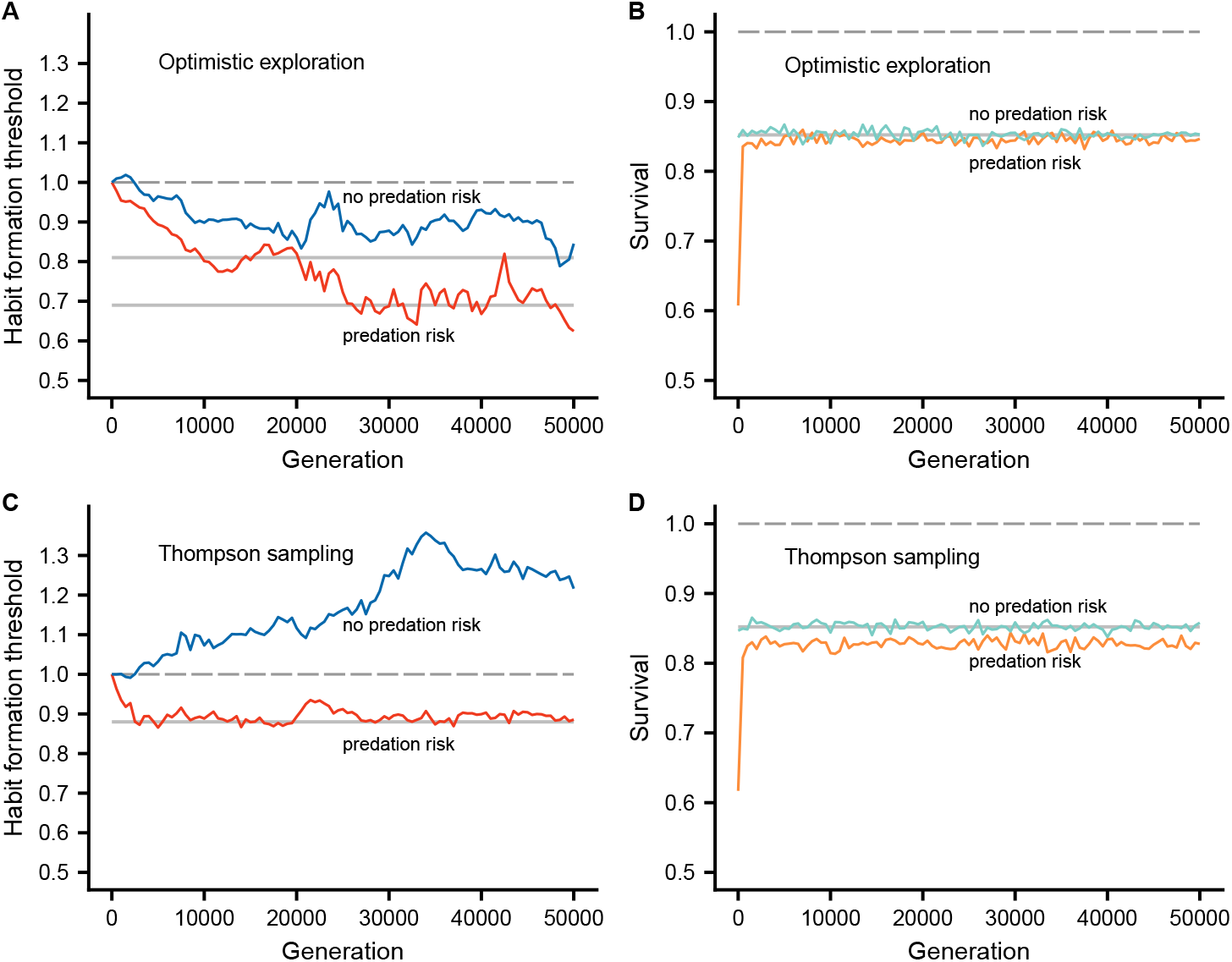
Illustration of evolutionary histories of habit formation thresholds *ĥ* and survival over 50,000 generations. Each panels shows a comparison between situations where habit formation influences predation risk (red/orange curves; *μ*_1_ = 0.00032) and where there is no influence (blue/green curves; *μ*_1_ = 0). The exploration method is OE in panels **(A)** and **(B)**, and TS in **(C)** and **(D)**. For OE, the parameters are as in as in Fig. 3C,D (*μ*_1_ = 0.00032) and Fig. 4C,D (*μ*_1_ = 0), and the trait values are kept fixed at these evolutionary equilibria, except that the trait *ĥ* is allowed to evolve from a starting point of *ĥ* = 1. The gray horizontal lines in **(A)** indicate the evolutionary equilibria for *ĥ*, and the line in **(B)** gives the theoretically expected survival of 0.85 without predation risk effects. For TS, parameters and traits are similarly as in Fig. S3C,D and Fig. S4C,D. For the predation risk cases, note that the survival rapidly increases from the starting value of 0.62 in both **(B)** and **(D)** (orange curves), showing that there is stronger selection to achieve rapid enough habit formation such that there is habitual foraging in most trials. Note also that the no-predation-risk threshold *ĥ* evolves to stay above *ĥ* = 1, so that habits do not form in **(C)**

### Parameters and trait values from simulations

**Table S1:** Trait values (mean over 100 simulations, each over 1000 generations) for 5 different cases of individual-based evolutionary simulations using OE and presented in the main text. The parameters that vary between cases are noise in rewards (*σ*_*R*_) and the influence of attention on predation risk (*μ*_1_), while *ρ*_0_ = 0, *σ*_0_ = 1, *μ*_e_ = 2, *σ*_e_ = 1, and two successive environments with 500 trials each hold for all cases.

| $\sigma_R, \mu_1 \times 10^3$ | $w_0$ | $\alpha_w$ | $\beta$ | $\alpha_h$ | $\hat{h}$ | $\hat{e}$ |
| --- | --- | --- | --- | --- | --- | --- |
| 0.25, 0.32 | 9.34 | 0.90 | 15.5 | 0.24 | 0.76 | 0.10 |
| 0.50, 0.32 | 5.76 | 0.64 | 12.6 | 0.15 | 0.76 | 0.10 |
| 0.10, 0.32 | 13.62 | 0.97 | 17.0 | 0.37 | 0.69 | 0.21 |
| 0.50, 0.0 | 4.62 | 0.62 | 15.8 | 0.06 | 0.88 | 0.07 |
| 0.10, 0.0 | 13.63 | 0.96 | 15.2 | 0.10 | 0.81 | 0.08 |

### Summary of notation for the models

**Table S2:**
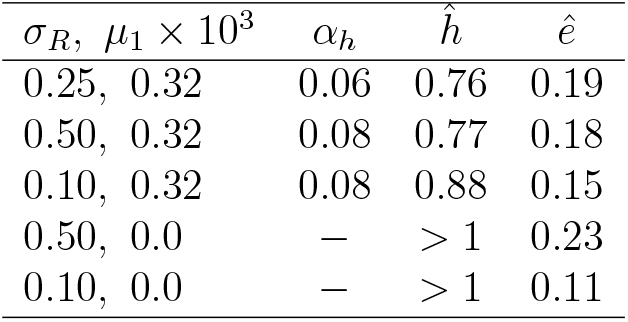
Trait values (mean over 100 simulations, each over 1000 generations) for 5 different cases of individual-based evolutionary simulations using TS and presented in the supplements. The parameters that vary between cases are noise in rewards (*σ*_*R*_) and the influence of attention on predation risk (*μ*_1_), while *ρ*_0_ = 0, *σ*_0_ = 1, *μ*_e_ = 2, *σ*_e_ = 1, and two successive environments with 500 trials each hold for all cases.

| $\sigma_R, \mu_1 \times 10^3$ | $\alpha_h$ | $\hat{h}$ | $\hat{e}$ |
| --- | --- | --- | --- |
| 0.25, 0.32 | 0.06 | 0.76 | 0.19 |
| 0.50, 0.32 | 0.08 | 0.77 | 0.18 |
| 0.10, 0.32 | 0.08 | 0.88 | 0.15 |
| 0.50, 0.0 | — | > 1 | 0.23 |
| 0.10, 0.0 | — | > 1 | 0.11 |

**Table S3:**
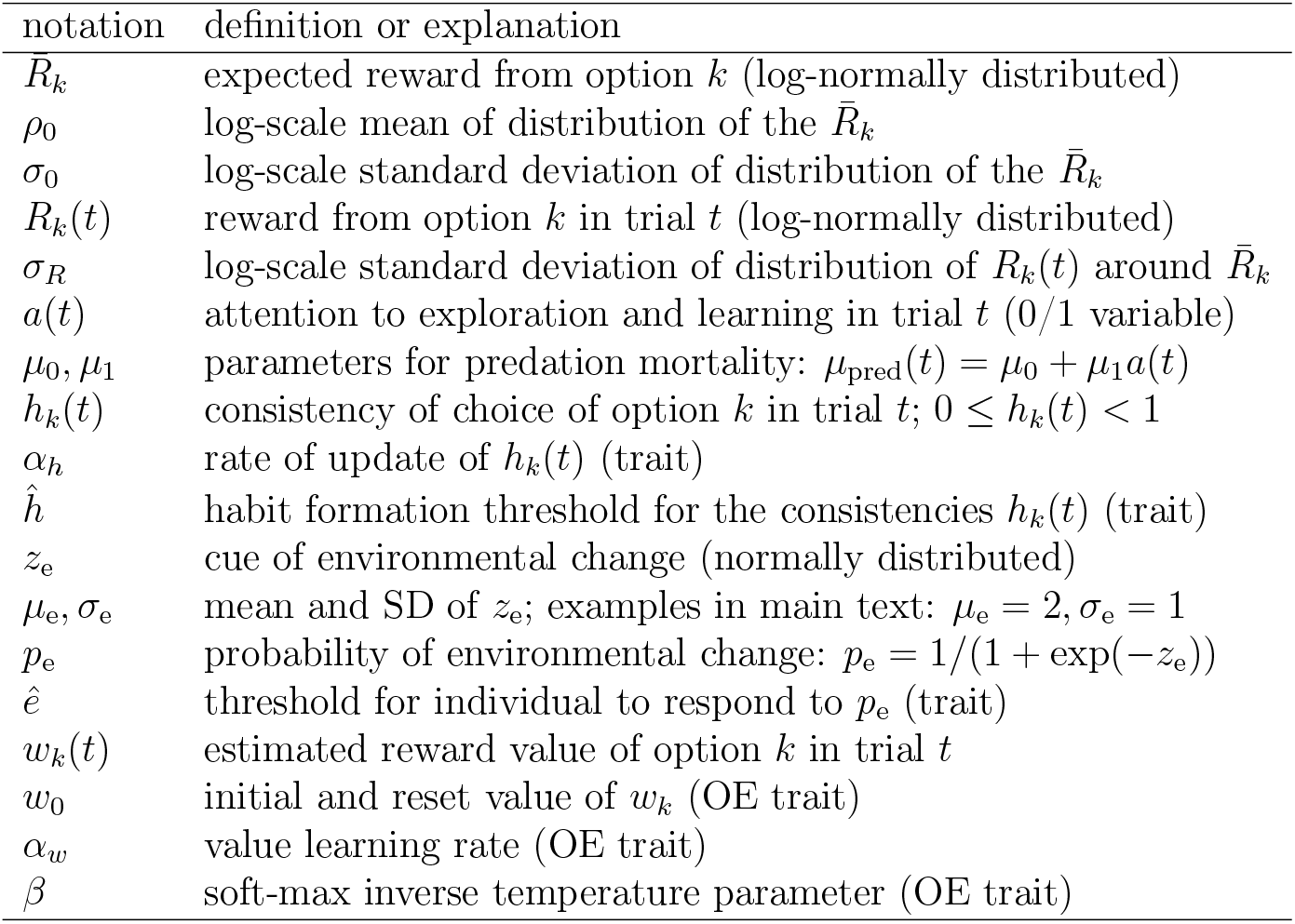
Table of notation for the OE and TS models.

| notation | definition or explanation |
| --- | --- |
| $\bar{R}_k$ | expected reward from option $k$ (log-normally distributed) |
| $\rho_0$ | log-scale mean of distribution of the $\bar{R}_k$ |
| $\sigma_0$ | log-scale standard deviation of distribution of the $\bar{R}_k$ |
| $R_k(t)$ | reward from option $k$ in trial $t$ (log-normally distributed) |
| $\sigma_R$ | log-scale standard deviation of distribution of $R_k(t)$ around $\bar{R}_k$ |
| $a(t)$ | attention to exploration and learning in trial $t$ (0/1 variable) |
| $\mu_0, \mu_1$ | parameters for predation mortality: $\mu_{\text{pred}}(t) = \mu_0 + \mu_1 a(t)$ |
| $h_k(t)$ | consistency of choice of option $k$ in trial $t$; $0 \leq h_k(t) < 1$ |
| $\alpha_h$ | rate of update of $h_k(t)$ (trait) |
| $\hat{h}$ | habit formation threshold for the consistencies $h_k(t)$ (trait) |
| $z_e$ | cue of environmental change (normally distributed) |
| $\mu_e, \sigma_e$ | mean and SD of $z_e$; examples in main text: $\mu_e = 2, \sigma_e = 1$ |
| $p_e$ | probability of environmental change: $p_e = 1/(1 + \exp(-z_e))$ |
| $\hat{e}$ | threshold for individual to respond to $p_e$ (trait) |
| $w_k(t)$ | estimated reward value of option $k$ in trial $t$ |
| $w_0$ | initial and reset value of $w_k$ (OE trait) |
| $\alpha_w$ | value learning rate (OE trait) |
| $\beta$ | soft-max inverse temperature parameter (OE trait) |

